# Structural basis for activation of glutaminase

**DOI:** 10.1101/2023.03.30.534948

**Authors:** Chen-Jun Guo, Zi-Xuan Wang, Ji-Long Liu

## Abstract

Glutaminase is a rate-limiting enzyme in glutaminolysis, which produces glutamate from glutamine and enters the TCA cycle[1]. In addition, it plays a key role in redox homeostasis[2], autophagy[3], immune system regulation[4], central nervous system maintenance[5], and senolysis[6]. Therefore, the allosteric regulation of glutaminase is a fascinating topic that has broad implications for our understanding of glutamine metabolism and related diseases[7–9]. Phosphate was discovered as a natural agonist for glutaminase in 1947[10], but the structural basis and mechanism for this regulation remains unclear. Using cryo-electron microscopy, here we determine the structure of human glutaminase with phosphate. This structure allows us to capture phosphate binding at the dimer-dimer interface at near atomic resolution, revealing an allosteric activation mechanism by remodelling the catalytic pocket. Surprisingly, we find that phosphate antagonizes BPTES (a classical antagonist) and CB-839 (the current subject of several phase II clinical trials). Accurate identification of phosphate binding sites lays the foundation for the design of glutaminase agonists and antagonists with broad pharmaceutical significance.

---

Glutaminase catalyzes the first step of glutamine metabolism, generates glutamate from glutamine, playing a critical role in various metabolic processes, including redox homeostasis, autophagy, immune system regulation, central nervous system maintenance and senolysis[2–9, 11–17]. In human, glutaminase is encoded by two different genes GLS1 and GLS2 [18]. GLS1 contains 19 introns and can be alternative spliced into two different active forms: kidney glutaminase A (KGA) and highly active glutaminase isoform C (GAC) [19] (Fig. 1A). GAC forms a tetramer when bound to substrate and can be further activated by anions such as phosphate (Pi) [10, 20, 21]. Additionally, it has been observed that GAC can form active filaments in vitro and in vivo[22–25] (Fig. 1B).

**Fig. 1.**
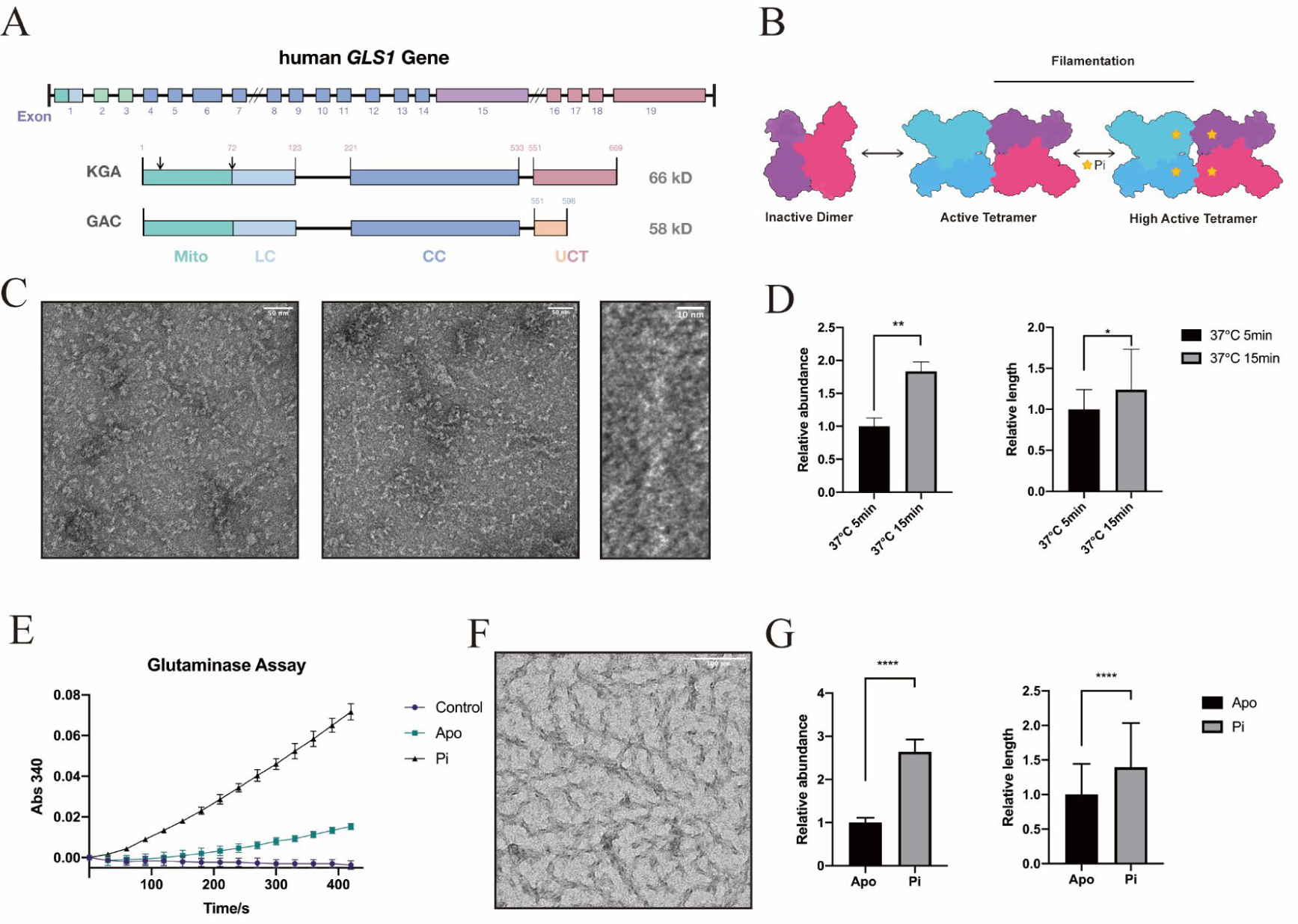
Domain organization and filamentation of glutaminase. **(A)** Domain organization of KGA and GAC isoforms of human GLS1 gene, showing their structural differences. Both KGA and GAC have a common N-terminal signal peptide and a Catalytic Core (CC) region. After mitochondrial transport, both forms lose the 72 amino acids in the head, leaving a low-complexity sequence from positions 72 to 123. The C-terminal domain of KGA consists of two ankyrin domains, while GAC has a unique C-tail without ankyrin domains. (Adapted from Charles J. McDonald, 2015) **(B)** Illustration of the oligomeric state and regulation mode of glutaminase. **(C)** Negative stain electron microscopy micrographs of GAC^APO^ incubated at 37 ℃ for 5 min and 15 min, revealing the time-dependent filamentation of GAC. The scale bar is defined separately in each graph. **(D)** Quantitative analysis of negative staining images from (C), indicating the stimulation of filamentation in GAC by incubation time. **(E)** Glutaminase assay for GAC^APO^ and GAC^Pi^ with control, demonstrating the activation of glutaminase by Pi. **(F)** Negative stain electron microscopy micrographs of GAC^Pi^ incubated at 37 ℃ for 10 min, showing the filamentation of glutaminase. The scale bar is defined in the graph as 100 nm. **(G)** Quantitative analysis of negative staining images for GAC^APO^ and GAC^Pi^, demonstrating the stimulation of glutaminase filamentation by Pi.

Metabolic reprogramming is a hallmark of cancer cells, known as the Warburg effect[26]. Glutamine, in addition to glucose, is another vital metabolite that provides a nitrogen source for amino acid synthesis and other biosynthetic pathways[16, 27]. Glutaminase, especially the GAC isoform, is highly overexpressed in various cancers for the utilization of glutamine[28–31]. Inhibition of glutaminase has shown promise in impeding cancer cell growth and inducing apoptosis, making it a compelling target for cancer therapy[7–9]. Currently, a specific glutaminase inhibitor is undergoing several clinical trials.

Interestingly, recent studies have demonstrated that the activation of glutaminase has a notable effect on eliminating cancer cell or solid tumors at the cellular or mouse levels, indicating that the regulation of glutaminase activation may have great potential for positive applications[32]. Additionally, the application value of glutaminase regulation has been verified in cellular or mouse experiments for processes such as non-alcoholic steatohepatitis[33], senescent cell clearance[6], and immune system reprogramming[4].

However, despite the progress made in understanding the role of glutaminase in cancer and other diseases, the activation and inhibition mechanisms of glutaminase remain enigmatic, limiting our understanding of its regulation and therapeutic potential.

By using Cryo-EM, we successfully resolved the structure of GAC bound with phosphate at a near atomic resolution. We reveal the binding site of the natural agonist phosphate and its remodeling effect on the catalytic pocket, ultimately elucidating the activation and inhibition mechanism of glutaminase. Additionally, our experiments and structural analyses have demonstrated that Pi and CB-839/BPTES act antagonistically towards each other. These findings not only enrich our understanding of the molecular basis for effective activation and inhibition, but also offer insights for the development of novel agonists and antagonists targeting glutaminase.

## Filamentation of GAC *in vitro* is a spontaneous and time-consuming process

GAC protein was expressed and purified using an N-terminal 6His SUMO expression system (Fig. S1). Negative staining samples were prepared and observed, which showed that GAC formed filamentous structures with a length of up to microns and a width of approximately 10 nm, resembling a crossed double helix (Fig. 1C).

The length of GAC filament increased with the incubation time. After incubation at 37°C for 5 minutes, the average length was approximately 120 nm and there were 8 filaments per micrograph. When the incubation time was extended to 15 min, the average length and number of filaments was 149 nm and 14.7 (Fig. 1D). Interestingly, when incubated on ice, few filaments formed (Fig. S2). Therefore, the formation of GAC filament is a spontaneous and time-consuming process which might be affected by temperature.

## Pi stimulates the reaction and filamentation

In 1935, Hans Kreb succeeded in isolating and characterizing three different forms of glutaminase that have distinct functions [1]. In 1966, it was discovered that both organic and inorganic acids can significantly enhance the activity of glutaminase, with Pi having the strongest effect [20, 21]. However, the precise mechanism by which acids promote glutaminase activity remains to be fully elucidated.

We discovered that purified GAC also has the characteristic of being activated by Pi. The presence of Pi resulted in a four-fold increase in the product of GAC, compared to the absence of Pi (Fig. 1E).

To investigate the effect of Pi on GAC, we prepared samples with phosphate and observed a significant increase in the degree of GAC filamentation under the same protein concentration, incubation time, and temperature (Fig. 1F). Through quantitative analysis, we found that Pi not only increased the number of GAC filaments but also increased the average length of GAC filaments (Fig. 1G).

Our observations suggest that Pi has the ability to promote both GAC catalysis and filamentation, implying the existence of a stable interaction between Pi and GAC.

## Filament structure of GAC-Pi

To investigate the interaction between Pi and GAC, as well as the assembly of GAC filaments, we prepared Cryo-EM samples containing Pi and an analog of glutamine, 6-Diazo-5-oxo-L-norleucine (DON). By using single-particle analysis processing and focus refinement strategy, we resolved the structure of the helical units and the helical interface of GAC filament at a resolution of 2.9 Å (Fig. S3 and S4; Table S1).

In the obtained GAC filament structure, GAC forms a tetramer as the helical unit, and the width of the entire filament structure is approximately 140 Å, with roughly 7 helical units in each cycle. The GAC filament has a helical twist of approximately 51°, and a rise of 70 Å (Fig. 2A). By viewing the filament from the top, using the characteristic N-termini of adjacent tetramers as a reference, the twist angle of approximately 51° can be observed (Fig. 2B).

**Fig. 2.**
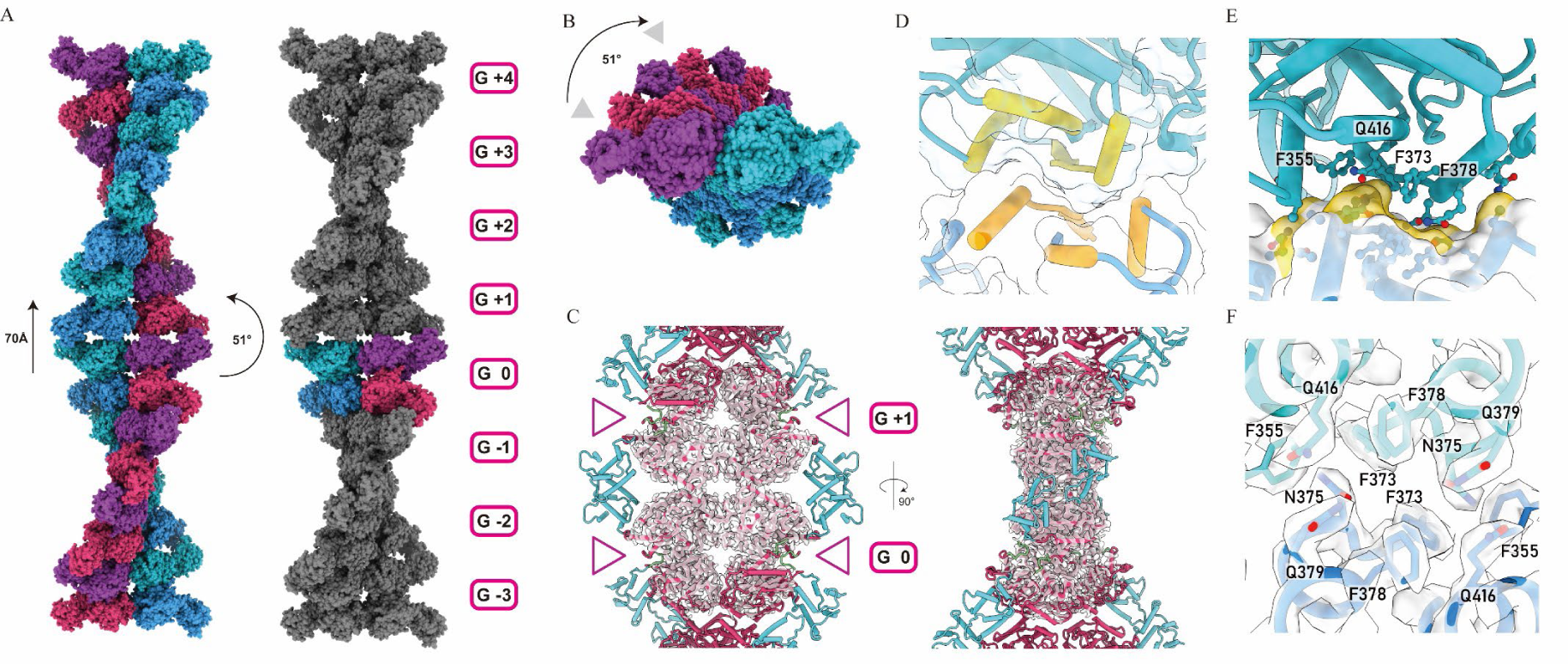
Filament structure of GAC-Pi. **(A)** GAC-Pi filaments assemble from GAC tetramers. Rise, twist, width and GAC tetramer extent are indicated. Left, filament colored by protomers; right, filament highlights one tetramer colored by protomers. **(B)** Top view of the GAC-Pi filament, demonstrating the helical twist in this orientation. **(C)** Front and side views of the helical interface of GAC-Pi filament, with map density revealing the arrangement of the catalytic core in the center of the filament, colored magenta, and N-termini and C-termini on the side, colored cyan and green, respectively. **(D)** Surface model of the helical interface, highlighting the secondary structures involved in filament formation. **(E)** Model and **(F)** map density of the helical interface residues, providing a detailed view of the interactions between GAC tetramers and the formation of the helical filament.

The N-termini and C-termini of GAC are situated on the side of the filament, with the catalytic core located on the rotational axis of the filament (Fig. 2C). The primary assembly interface of the filament is concentrated on the catalytic core, while there may also be some interactions in the N-terminal structural domain on both sides (Fig. S5).

The assembly interface of the GAC filament is symmetrical, owing to the helical unit’s symmetry. It comprises two identical parts that two adjacent protomers complement each other in a flip-flop manner. The secondary structures involved in the assembly interface are α helices 351-365, 375-384, 407-419 and β barrel 371-373 (Fig. 2D).

Focusing on one of the interaction interfaces, it shapes like a narrow rectangle, with a total interface area about 585 Å^2^ (Fig. 2E). Sequence alignment shows that the amino acids involved in the helical interface, F355, F373, N375, F378, Q379, and Q416, are highly conserved in mammalian GLS, including humans, mice, and rats (Fig. S6).

Through computational analysis, we found that Q416 forms a salt bridge interaction with N375 in the other protomer, contributing to the force that drives filament formation (Fig. 2, E and F). F355, F373, and F378 are arranged around Q416 in an enveloping manner. F355 forms a cation-pi interaction with N375 in the other protomer, contributing to another force that drives filament formation (Fig. 2, E and F).

## Pi binding site locates on the dimer-dimer interface of GAC

Since the discovery that anions could activate GAC, people have been investigating the binding sites and the regulation modes of anions on GAC. Among anions, phosphate has received significant attention as it is a vital inorganic acid in mitochondrial and is known to enhance GAC activity considerably. Structures of GAC with different ligands have been resolved [18, 34–44]. However, the binding mode and mechanism of phosphate to activate GAC remain elusive.

By using the electron density maps with near-atomic resolution, we were able to observe the conformation of the GAC tetramer in the filament and identify the allosteric binding site of Pi in GAC (Fig. 3A). GAC tetramer has two different assembly interfaces among protomers: the dimer interface and the dimer-dimer interface.

**Fig. 3.**
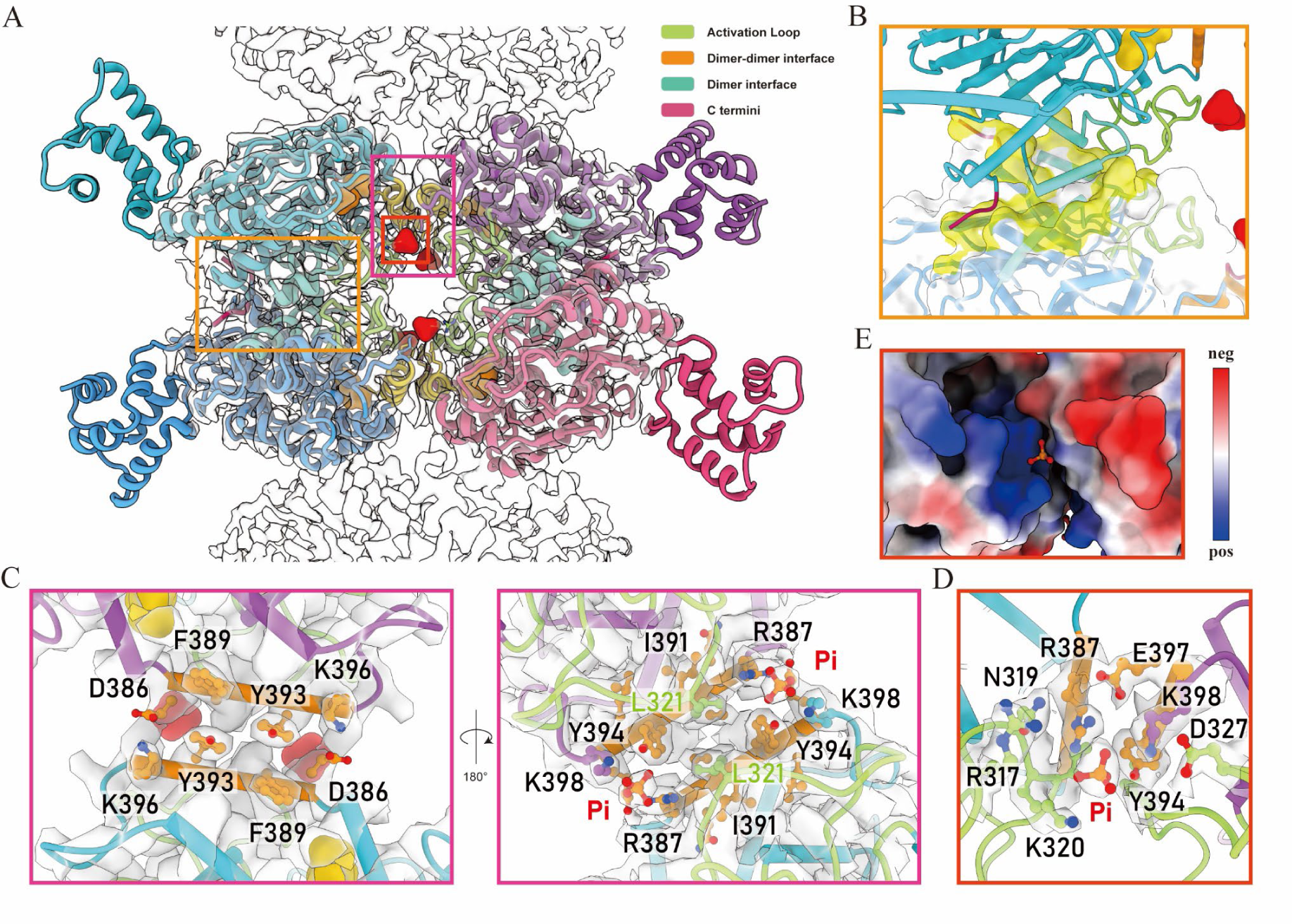
Pi binding site locates on the dimer-dimer interface of GAC. **(A)** Cryo-EM structure of one helical unit of GAC-Pi filament, with different regions of the protein colored. The Pi binding site, located on the dimer-dimer interface, is indicated with the surface of Pi colored in red. **(B)** Zoom-in view of the dimer interface, showing the contacting area colored in yellow in the surface model. **(C)** Model and density map of the dimer-dimer interface, demonstrating the interactions between Pi, L321 from AL with helix 386-397. **(D)** Model and density map for the binding site of Pi on the dimer-dimer interface, revealing the clear density of Pi and composition of the binding pocket.

The dimer interface is mainly composed of the activation loop 308-334, helices 450-463, 469-475, loop 528-532, and partial C tail (Fig. 3B). It is highly conserved from prokaryotes to eukaryotes and is necessary for the formation of glutaminase dimers. The dimer-dimer interface is composed of interactions between a pair of reverse parallel helices, specifically helix 386-397, from two protomers. The pi-pi interaction between F389 and Y393 and the electrostatic interaction between D386 and K396 in different protomers were believed to play a critical role in stabilizing this interface.

In addition, in the presence of phosphate binding, the activation loop also participates in stabilizing this interface. L321 in Loop 308-334 is embedded in helix 386-397, surrounded by Y394, I391 and R387, increasing the contact area of the dimer-dimer interface (Fig. 3C).

A GAC tetramer contains four phosphate binding sites, which are located on the dimer-dimer interface consisting of R317 and K320 on the activation loop, as well as R387 on the dimer-dimer interface of one protomer and Y394 and K398 on the dimer-dimer interface of another protomer (Fig. 3, A and D). Sequence alignment reveals that these amino acids involved in phosphate binding are highly conserved in mammalian GAC (Fig. S6). The binding site is characterized by multiple positively charged amino acids that interact with phosphate, resulting in a strong overall positive charge. This structural feature provides a basis for the requirement of multivalent acids to activate GAC activity fully (Fig. 3E). During data processing, we observed multiple binding modes of phosphate ions in different tetramers and different locations of the same tetramer through 3D classification, indicating the relative flexibility of the phosphate binding site (Fig. S7).

## Filamentation by Pi remodels the catalytic pocket

Significant conformational changes were observed in the GAC tetramer when comparing the tetramers in filament to free tetramers. To avoid confusion, we will use the term "GAC-PF tetramer" to refer specifically to the filamentous GAC tetramer bound with phosphate.

First, the overall conformation of the GAC-PF tetramer has changed comparing to the free tetramer (Fig. 4A). Specifically, the GAC-PF tetramer is compressed along the filament formation axis, resulting in an inward rotation of the four catalytic cores and a relative outward expansion of the N-terminal domain comparing to the free tetramer. The tetramer interface also experiences slipping and alterations, bringing the four catalytic sites closer together, while the outward expansion of the N-terminal domain creates more space for the C-tail.

**Fig. 4.**
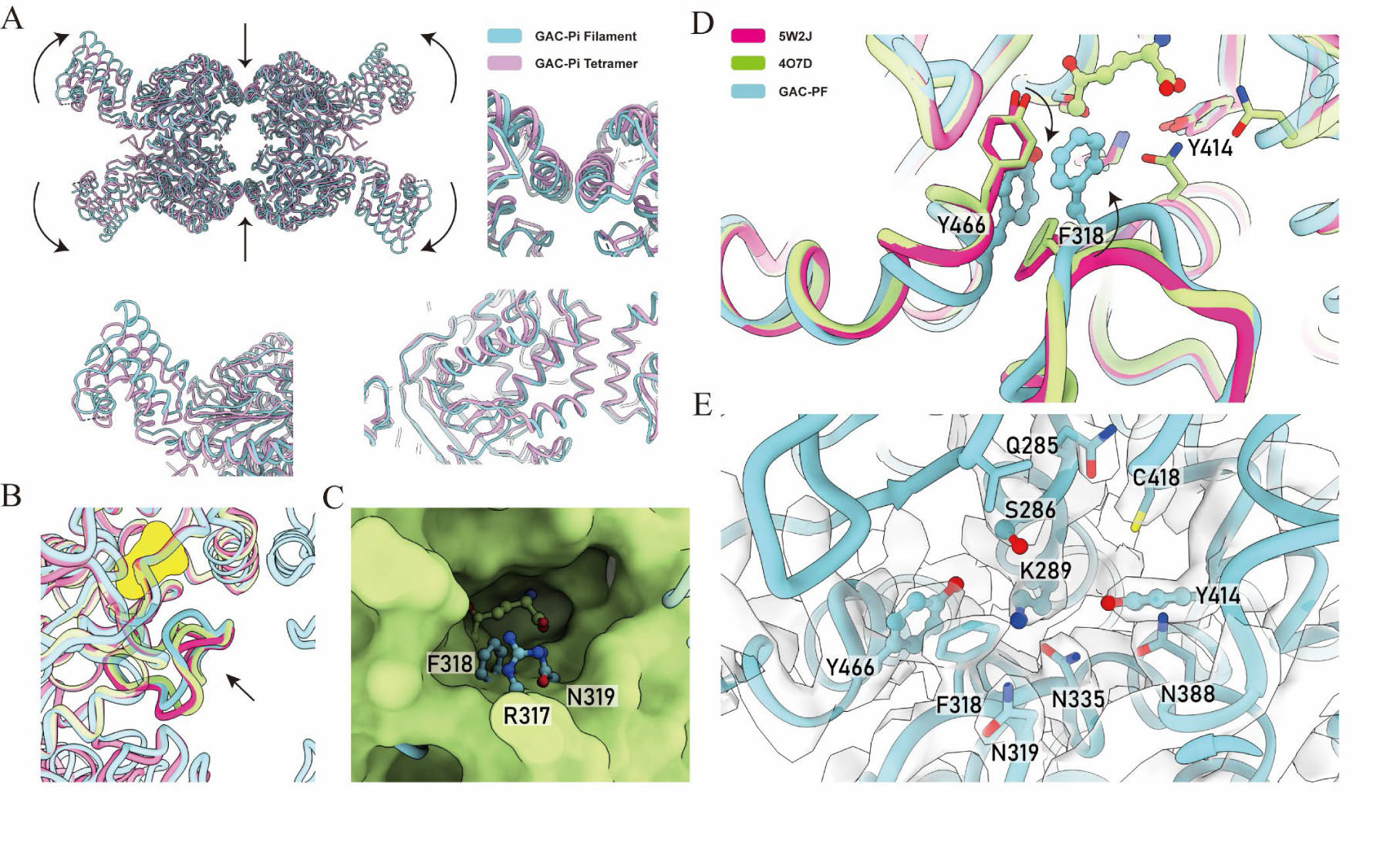
Filamentation by Pi remodels the catalytic pocket of glutaminase. **(A)** Comparison of the GAC-PF tetramer and GAC-Pi tetramer models reveals significant conformational changes. **(B)** The active loop (AL) region in GAC-PF tetramer is reshaped towards the catalytic center which is displayed by yellow surface, compared to AL obtained in other models (PDB code: 4O7D and 5W2J). **(C)** The reshaped AL further fills the catalytic pocket, as evidenced by the green surface of the 5W2J model being filled with residues R317, F318, and N319 from the AL of GAC-PF. **(D)** The Y466 loop undergoes a conformational change in the GAC-PF tetramer with the interaction of the reshaped AL. **(E)** The detailed composition of the remodeled catalytic pocket of glutaminase is shown with the density map.

Second, the activation loops of each protomer in the GAC-PF tetramer have been reshaped and stabilized (Fig. 4B). The activation loop (AL, residues 308-334) is a long loop that is thought to be involved in the activation of the GLS reaction and is highly dynamic, being absent in most GAC structures (Fig. S8). However, in the GAC-PF tetramer, the B-factor analysis of our models shows that the activation loop is in a stable state (Fig. S9). Compared to the AL obtained from glutaminase tetramer in DON-inhibited state and the dimer formed by disrupting the dimer-dimer interface through D386K mutation (PDB code: 4O7D and 5W2J), the activation loop bound to phosphate exhibits a more closed conformation towards catalytic core (Fig. 4B).

The closed AL alters the catalytic pocket of GAC. First, the stability of the AL further fills the catalytic pocket (Fig. 4C). Secondly, the pi-pi interaction between F318 and Y466 changes the position of the Y466 loop, reducing the steric hindrance between Y466 and F318 and increasing the affinity of the pocket (Fig. 4D). This explains the decrease in enzyme activity caused by F318A reported by J. Sivaraman in 2012 [24]. Finally, the downward movement of Y466, N335, and the activation loop compared to the unbound state also reshapes the reaction pocket (Fig. 4E).

During the catalytic process, Y414 and Y466 act as proton transferors, while K289 acts as a proton donor [45] (Fig. S10). In the remodeled catalytic pocket, Y466, K289, and Y414 are located on the same plane, and the distance between Y466 and K289 is closer, which will facilitate proton transfer and product release, thereby accelerating the reaction (Fig. 4E).

Our structure provides a precise view of the conformational changes and the remodeling of the catalytic pocket of GAC induced by the binding of phosphate and filamentation. This serves as a structural basis for understanding of how GAC is activated by phosphate.

## Pi antagonizes BPTES/CB-839, two classical inhibitors of glutaminase

BPTES is a non-competitive inhibitor targeting GLS1, which was developed in 2008 and demonstrated efficient and precise inhibition (Fig. 5A). BPTES binds to the dimer-dimer interface and interacts with two different protomers, with two different binding sites for BPTES in one tetramer. Based on this, subsequent research has developed the more effective CB-839 (Telaglenastat) (Fig. 5A). It can not only more strongly inhibit the two active isoforms of GLS1, but also induce cell apoptosis and has powerful anti-tumor activity. Telaglenastat is currently still in a series of clinical trials. Although both BPTES and CB-839 have fascinating specificity and anti-tumor activity, the mechanism underlying glutaminase inhibition by these inhibitors remains unclear.

**Fig. 5.**
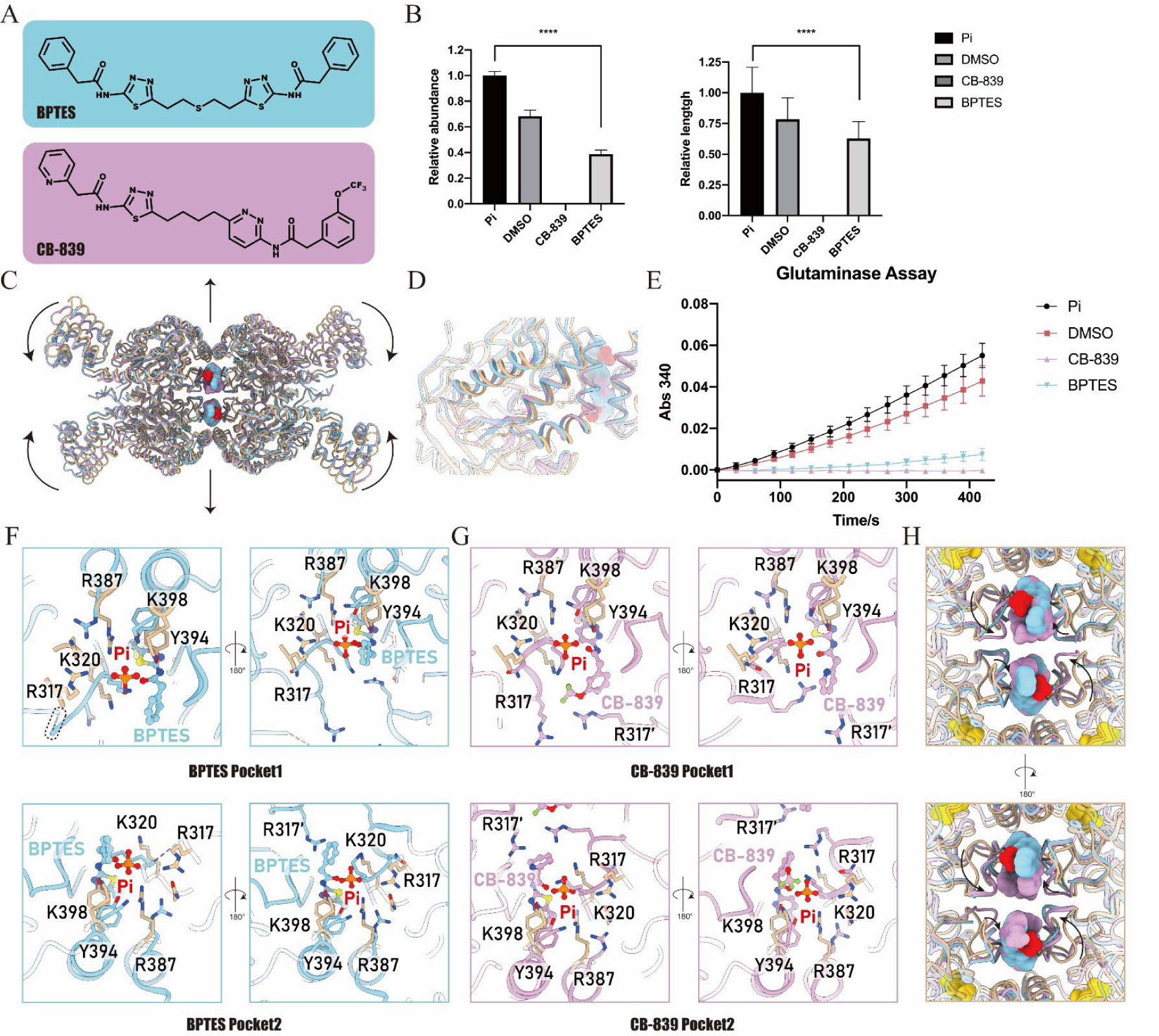
Pi antagonizes BPTES/CB-839, two classical inhibitors of glutaminase. **(A)** Chemical structures of BPTES and CB-839, two representative inhibitors of glutaminase. **(B)** Quantification of negative staining images showing the disruption of GAC filamentation by BPTES and CB-839. **(C)** Overall conformational changes of the GAC tetramer induced by BPTES or CB-839 binding. The GAC PF model is colored in champagne, while the BPTES or CB-839 bound model is colored in cyan or pink, respectively. **(D)** Conformational changes of the helical interface upon binding of BPTES or CB-839. **(E)** Inhibition of GAC by BPTES or CB-839 shown by glutaminase assay for GAC^Pi^ with or without inhibitors, and DMSO as a control. **(F, G)** Detailed conformational comparison of GAC-PF with BPTES or CB-839 bound GAC tetramer, illustrating the conformational changes of the binding site and the dimer-dimer interface. Antagonistic binding of Pi and inhibitors is shown. The disordered loop is indicated by a dashed box. **(H)** Detailed conformational changes of AL upon binding of BPTES or CB-839, highlighting the interaction between the inhibitors and AL. The antagonistic interaction mode of Pi and inhibitors with AL is shown. The coincide of Pi and inhibitor in the binding site is also depicted.

We aimed to examine the impact of inhibitors on GAC filament formation by adding BPTES and CB-839 to pre-existing filament samples. Initially, GAC was subjected to incubation at 37°C in a 40 mM phosphate system for 10 minutes, resulting in the formation of GAC filaments in the system. Upon the addition of both inhibitors, they demonstrated varying degrees of efficacy in disrupting the formation of the filaments and converting GAC into free tetramers (Fig. S11). Our quantitative analysis revealed that BPTES significantly reduced the abundance of GAC filaments and shortened the length of each filament. In contrast, CB-839 exhibited stronger ability in disassembling GAC filament and no filamentous structure was observed in the quantitatively counted images. Instead, the main form of GAC was transformed into a tetramer (Fig. 5B). These findings suggest that both inhibitors may cause disassembly of the filamentous structure by altering the conformation of the tetramer.

The conformational changes of the tetramer explain the effect of inhibitors on GAC filament formation (Fig. 5, C and D). When BPTES or CB-839 bound to GAC, the activation loop interacted with the inhibitor, causing a change in the dimer-dimer interface. This led to the external rotation of the GAC catalytic core and the internal shrinkage of the N-terminal, as compared to the GAC-PF tetramer. The inhibited tetramer exhibited a mode of expansion compared with GAC-PF (Fig. 5C). In addition, the active interface of the filament also exhibited slipping and changes, which hindered the inhibited tetramer from forming the filamentous structure in the mode of GAC-PF (Fig. 5D).

Glutaminase activity experiments conducted simultaneously with NS-EM demonstrated varying degrees of inhibition of GAC activity by BPTES and CB-839. Specifically, at a concentration of 0.9 μm, BPTES exhibited significant inhibition of GAC’s reaction, while CB-839 almost completely inhibited GAC’s activity (Fig. 5E).

Comparing GAC-PF tetramer with inhibitors bounded tetramer, we found that the natural activator Pi and inhibitors are antagonistic competitive ligands of GAC:

On the one hand, both Pi and inhibitors bind to the dimer-dimer interface. When two tetramers align, the binding of inhibitors and Pi overlaps with each other, which creates steric hindrance and directly limits Pi binding and activation (Fig. 5C and 5H). The overlap of the binding sites also suggests that Pi and inhibitors cannot coexist in GAC.

Furthermore, when inhibitors bind, the entire Pi binding site undergoes reshaping, preventing Pi from binding to GAC in the same manner as GAC-PF (Fig. 5, F and G). Conversely, when Pi is bound to GAC, the binding of inhibitors is also hindered due to the overlap of the binding sites (Fig. 5, F and G). This provides a direct structural basis for the mutual antagonism of inhibitors and the natural activator Pi.

On the other hand, compared to the ordered fixation of the activation loop by Pi, inhibitor binding fixes the activation loop in a different mode, keeping it away from the catalytic active site, making the catalytic reaction less likely to occur (Fig. 5, F and H) (Fig. S12). The binding of BPTES still leaves an active area for the loop, and the B-factor also shows that the end of BPTES may still be in a relatively flexible state, allowing for some movement in the loop. CB-839 exhibits more potent inhibition, as the entire activation loop is fixed, and the B-factor shows that the binding of CB-839 is also very stable, making AL more fixed and therefore more strongly inhibited (Fig. 5, G and H) (Fig. S12).

## Discussion

GLS plays multiple important roles in various biological processes, making its allosteric regulation of great potential value. Currently, cancer-related applications receive the most attention in research and development[7, 9, 12, 13, 24, 31, 46, 47]. Specific inhibitor CB-839 is the most extensively studied in clinical trials, but its monotherapy effect is not as successful as expected[46, 48]. Combinatorial therapeutic strategies with other treatments are still under investigation, although the combined therapies with Everolimus and Cabozantinib are not significant[49, 50]. However, the clinical use of CB-839 in combination with radiotherapy has shown positive results[51].

Full antagonists of GLS may not directly achieve immediate effects in cancer treatment and bring about some side effects. This may be related to the key roles of glutaminase in various biological processes, such as glutamine deprivation-induced apoptosis[32] and immune cell proliferation and differentiation[52].

Under these circumstances, partial agonist-antagonists with a ceiling effect provide a new option. Their use could achieve the maximum inhibition of abnormal glutaminolysis while minimizing the impact on normal cells.

Besides, the safety demand is particularly important for the implications in other therapies such as non-alcoholic steatohepatitis, regulating immune cell differentiation, and clearing senescent cells.

Recently, in cell experiments and mouse models, promoting the highly active GLS1 filament formation suppresses glutamine addicted cancer cell growth and tumorigenesis[32]. Therefore, the design of full agonists for glutaminase has been proposed as the next potential therapeutic avenue for cancer.

Our work has revealed the long-sought-for activation mechanism, completed the inhibition mechanism of glutaminase, and enhanced our understanding of glutaminase regulation. The identification of allosteric activation sites and filamentous structures provides a structural basis for the design of various potential antagonists, partial agonist-antagonists, and complete agonists with potential pharmaceutical value.

## Methods

### GAC plasmid construction and protein purification

The human GAC (amino acids 123-598) gene was cloned into pET28a vector with a N-terminal 6 × His SUMO[53] tag and transformed into Escherichia coli Transetta (DE3) cells for expression. Transformed cells were cultured in LB medium at 37 °C with 220 rpm until OD600 reached a range of 0.4 to 0.6,and then down-tempered to 18 °C for 0.5 to 1 hr before induction with 0.1 mM IPTG for 16-20 hr at 18 °C. Cells were pelleted by centrifugation at 4,500 rpm for 18 min followed by resuspension in cold lysis buffer (50 mM Tris-HCl pH 8.5, 500 mM NaCl, 10% glycerol, 20mM imidazole, 1 mM PMSF, 5mM β-mercaptoethanol, 5mM benzamidine, 2 μg/ml leupeptin, and 2 μg/ml pepstatin). Bacteria in lysis buffer were disrupted by high pressure at 750 bar and centrifuged at 18,000 rpm for 1 hr at 4 °C to collect supernatant. This was incubated with equilibrated Ni-Agarose (Qiagen) for 1 hr. Next, Ni-Agarose was washed by washing buffer (50 mM Tris-HCl pH 8.5, 500 mM NaCl, 10% glycerol, 40mM imidazole), and proteins were eluted with elution buffer (30 mM Tris-HCl pH 8.5, 100 mM NaCl, 240 mM imidazole,5 mM β-mercaptoethanol), peak fractions were treated with SUMO protease ULP1 for 1 hr at 8 °C. SuperoseTM 6 Increase 10/30 GL column and AKTA Pure (Cytiva) were used for further purification. Finally, GAC was eluted with buffer containing 100 mM NaCl and 30 mM Tris-HCl pH 8.5.

### Glutaminase Assay

GAC activity was determined using the L-Glutamate Dehydrogenase-based (GDH), two-step Glutaminase protocol as previously published[54]. GAC was incubated in the reaction buffer containing 30 mM Tris-HCl pH 8.5, 10 units GDH and 2 mM NAD+ for 5-10 min at room temperature or 37 °C. For inhibitor assay, 0.9 μM inhibitors, which were dissolved by DMSO, were added to the system, and then incubated at 37 °C for 10 min. To initiate reaction, 50 mM glutamine was added into the mixture. Absorption of a wavelength of 340 nm of each reaction mixture was measured with SpectraMax i3 as the indication for NADH levels at individual time points and absorbance represents the glutaminase activity.

### Negative Staining

13 μM GAC was incubated for 10 min at 37 °C for preparation of GACApo sample. In other cases, GAC proteins were first incubated with 40 mM Pi for 10 min at 37 °C, and then mixed with inhibitors with concentration of 0.9 μM for 10 min at 37 °C. The prepared protein samples were applied to glow-discharged carbon-coated EM grids (400 mech, EMCN), and stained with 1% uranyl formate. Images were acquired at 57,000× magnification using a Tecnai Spirit G21 microscope (FEI).

### Cryo-EM Grid Preparation and Data Collection

For preparing the Pi-bound GAC sample, 12 μM GAC was first incubated with 2.3 mM DON for 30 min at 37 °C, and then incubated with 40 mM Pi for 5 min at 37 °C. Samples were prepared with 100 holey carbon film (Q46208-Cu200-R0.6/1) and FEI Vitrobot (4 °C temperature, 3.5 s blotting time, −1 blot force). Images were taken with a Gatan K3 summit camera on a FEI Titan Krios electron microscope operated at 300 kV. The magnification was 22,500 × in superresolution mode with the defocus rage −1.2 to −1.8 μm and a pixel size of 1.06 Å. The total dose was 50e−/Å2 subdivided into 40 frames at 2.8-s exposure using SerialEM.

### Image Processing

The whole workflow was done in RELION 3.1.2. Raw movies were dose weighted and aligned by MOTIONCOR2 through RELION3 GUI, and contrast transfer function (CTF) parameters were determined by CTFFIND4. 3,271,231 particles were picked by autopicking. After two-dimensional and three-dimensional (3D) classification with C1 and D2 symmetry, 611,365 particles were selected for the 3D refinement. 321,414 particles centering on interface generated a map of 3.0 Å, and 203,004 particles centering on tetramer generated a map of 3.1 Å. CTF refinement and Bayesian polishing were applied to each particle. The 3D refinement and continued focus refinement with tight mask were used in two different particle sets. Finally, we constructed maps with resolution of 2.9 Å for GAC-Pi filament both centering on the helical interface and helical unit.

### Model Building and Refinement

Previous model [Protein Data Bank (PDB) ID:3SS4] was applied for the initial model. Model of monomer was manually refined according to the electron density map with Coot software[55]. The refined monomer model was symmetrized to build tetramer models in Chimera software[56]. The tetramer models were subsequently real-space refined in Python-based hierarchical environment for integrated xtallography (Phenix) software[57].

### Quantification and statistical analysis

Images of negative staining is processed using ImageJ[58] for quantification. Polymers of length greater than 50 nm were considered as filaments. Results of quantification and glutaminase assay were analyzed using GraphPad Prism 8[59] and were shown as means ± SD of three or more independent experiments. Mafft program[60] was used for alignment of sequence of human KGA and GAC (UniProtKB:O94925;homo sapiens), human GLSL (UniProtKB:Q9UI32;homo sapiens), rat GLSK (UniProtKB:P13264;Rattus norvegicus), rat GLSL (UniProtKB:P28492;Rattus norvegicus), mouse GLSK (UniProtKB:D3Z7P3;Mus musculus), mouse GLSL (UniProtKB:Q571F8;Mus musculus), lion GLS (UniProtKB:A0A8C8XW40,Panthera leo), pig GLS (UniProtKB:A0A4X1THT5,Sus scrofa), and goat GLS (UniProtKB:A0A452DX27,Capra hircus). The result of sequence alignment was visualized by ESPript 3[61] which rendered sequence similarities and structure information taking crystal structure of dimeric form of mouse Glutaminase C (PDB EntryID:5W2J) as reference.

## Data availability

The structure data accession codes are EMD-35573, EMD-35574, and PDB-8IMA, PDB-8IMB.

## Acknowledgments

We thank Suwen Zhao for her helpful discussions. EM data were collected at the ShanghaiTech Cryo-EM Imaging Facility. We thank the Molecular and Cell Biology Core Facility (MCBCF) at the School of Life Science and Technology, ShanghaiTech University and Shanghai Frontiers Science Center for Biomacromolecules and Precision Medicine for providing technical support. This work was supported by grants from: Ministry of Science and Technology of China (No. 2021YFA0804700), National Natural Science Foundation of China (No. 31771490), Shanghai Science and Technology Commission (No. 20JC1410500), UK Medical Research Council (grant nos. MC_UU_12021/3 and MC_U137788471) for grants to J.L.L.

## Author contributions

Conceptualization: C.J.G., J.L.L. Methodology: C.J.G. Investigation: C.J.G., Z.X.W.

Visualization: C.J.G. Funding acquisition: J.L.L.

Project administration: C.J.G., J.L.L. Supervision: C.J.G., J.L.L.

Writing – original draft: C.J.G.

Writing – review & editing: C.J.G., Z.X.W., J.L.L.

## Supplementary information

**Fig. S1.**
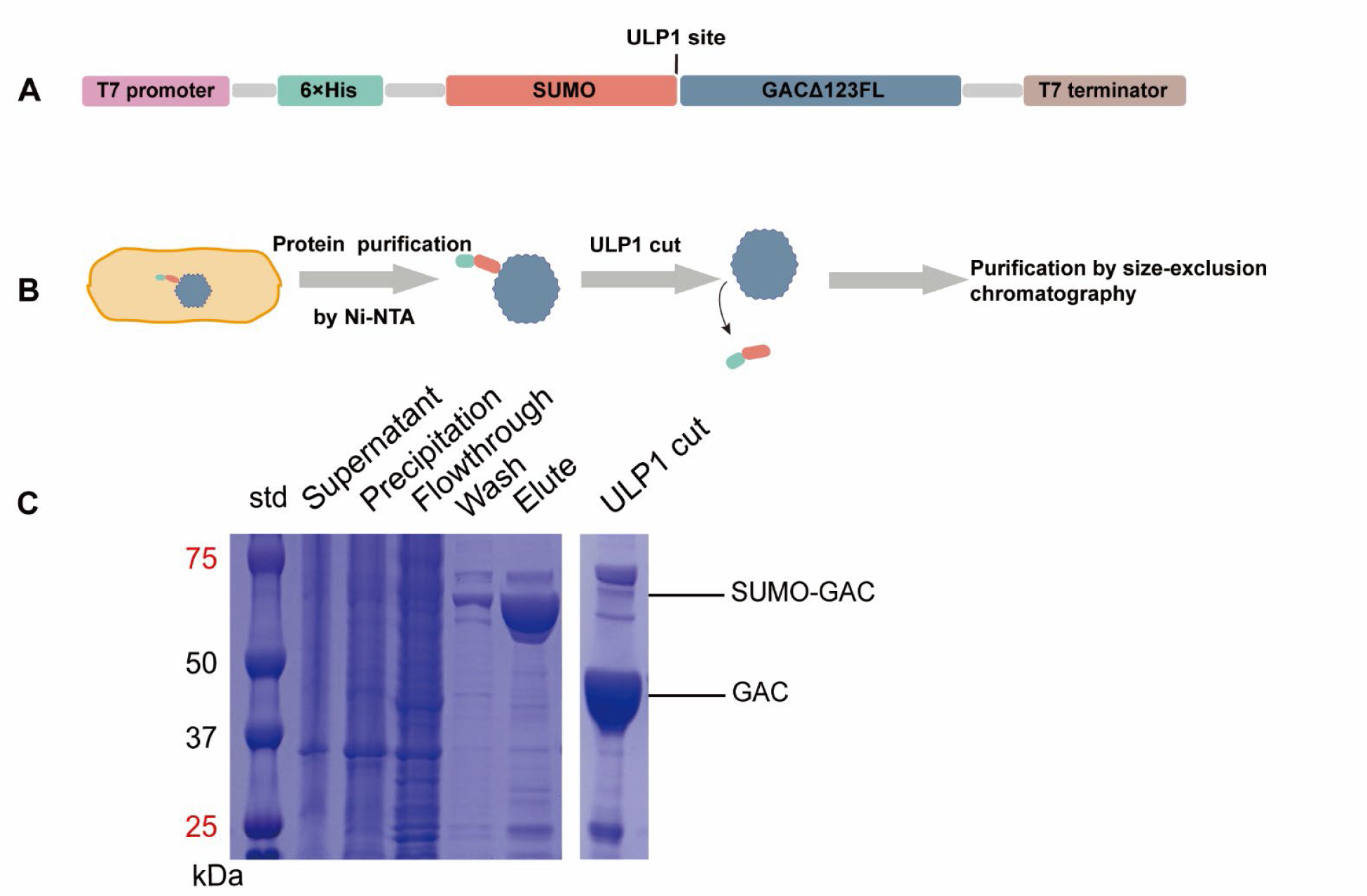
GAC plasmid construction and protein purification. A) Plasmid construction. The plasmid was designed for the expression of hGAC with a 6×His SUMO tag at the N-terminus. B) Protein purification of hGAC by Ni-NTA. The 6×His SUMO tag was cleaved by ULP1, followed by further purification by size-exclusion chromatography. C) SDS-Page analysis of purified hGAC proteins showing the cleavage of SUMO tag.

**Fig. S2.**
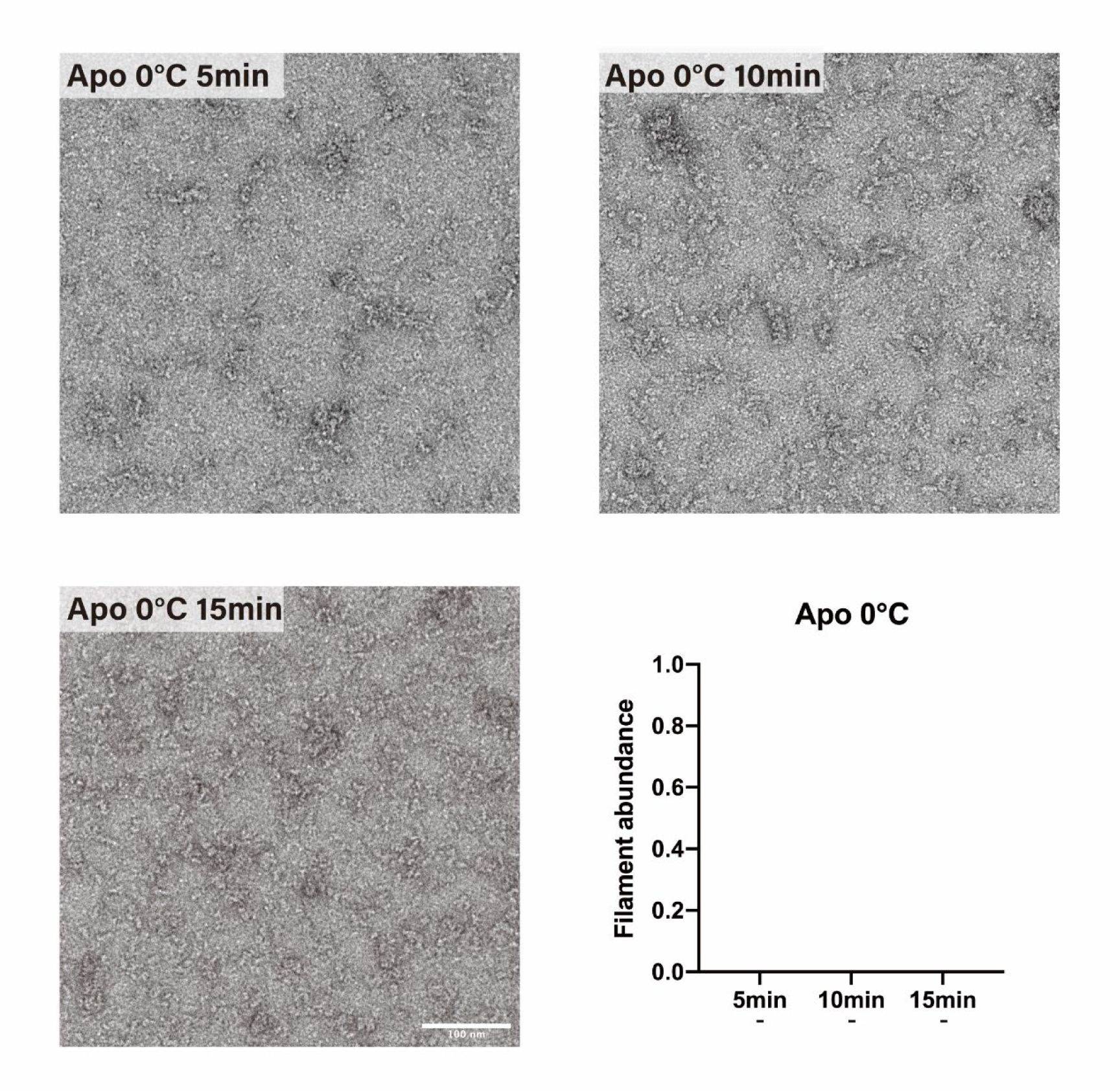
Few filaments formed on ice. Negative stain electron microscopy micrographs of GAC^APO^ incubated at 0 ℃ for 5,10 and 15min are shown above. The scale bar is defined in the graph as 100 nm. The quantification of filament abundance is also presented, where "-" indicates that no filament was found in the image.

**Fig. S3.**
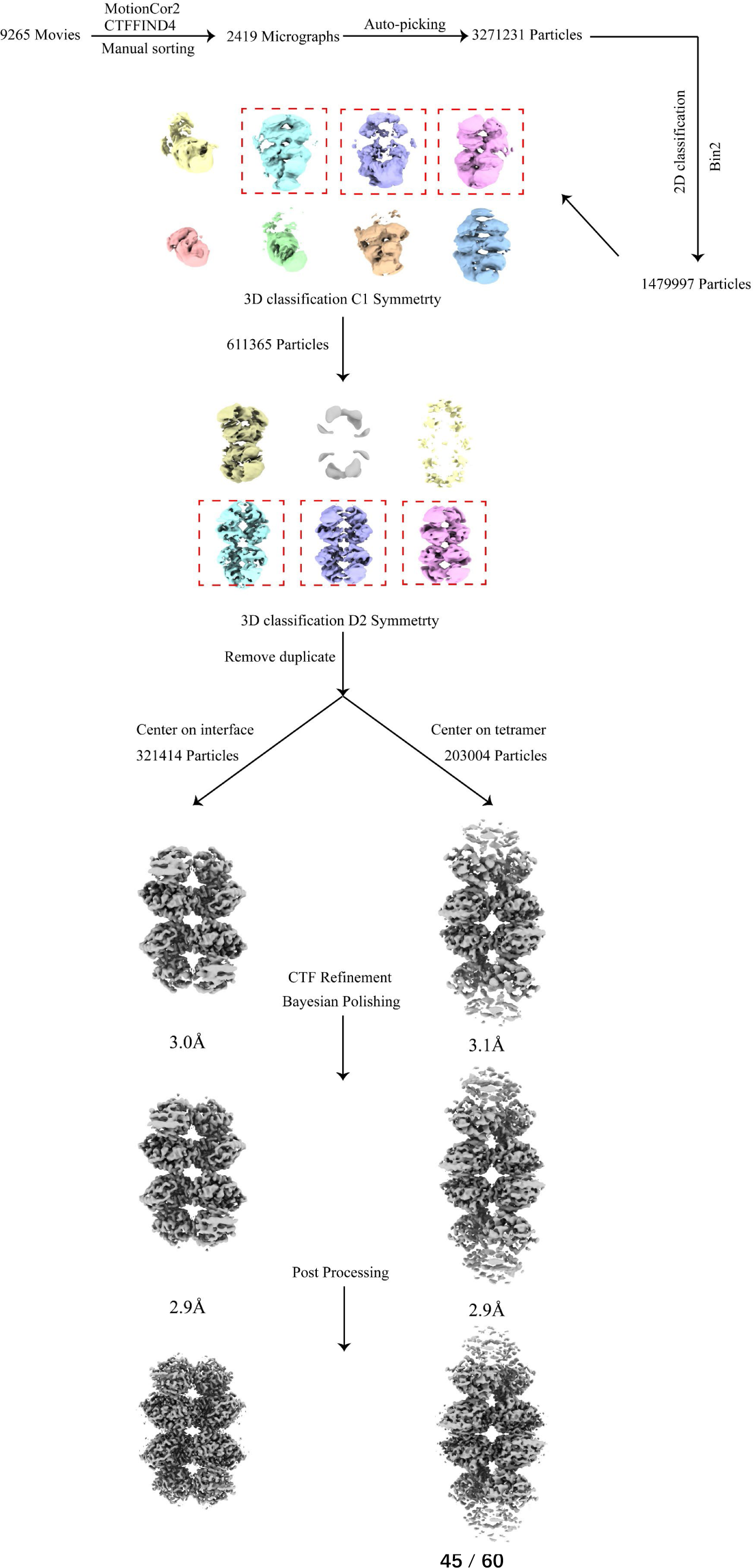
The workflow of data processing. After initial particle picking and 2D classification, 3D classification was performed to select well-aligned filaments. Focused refinement was then carried out on both the helical unit and helical interface to improve the resolution of the final structures. Further details can be found in the Method section.

**Fig. S4.**
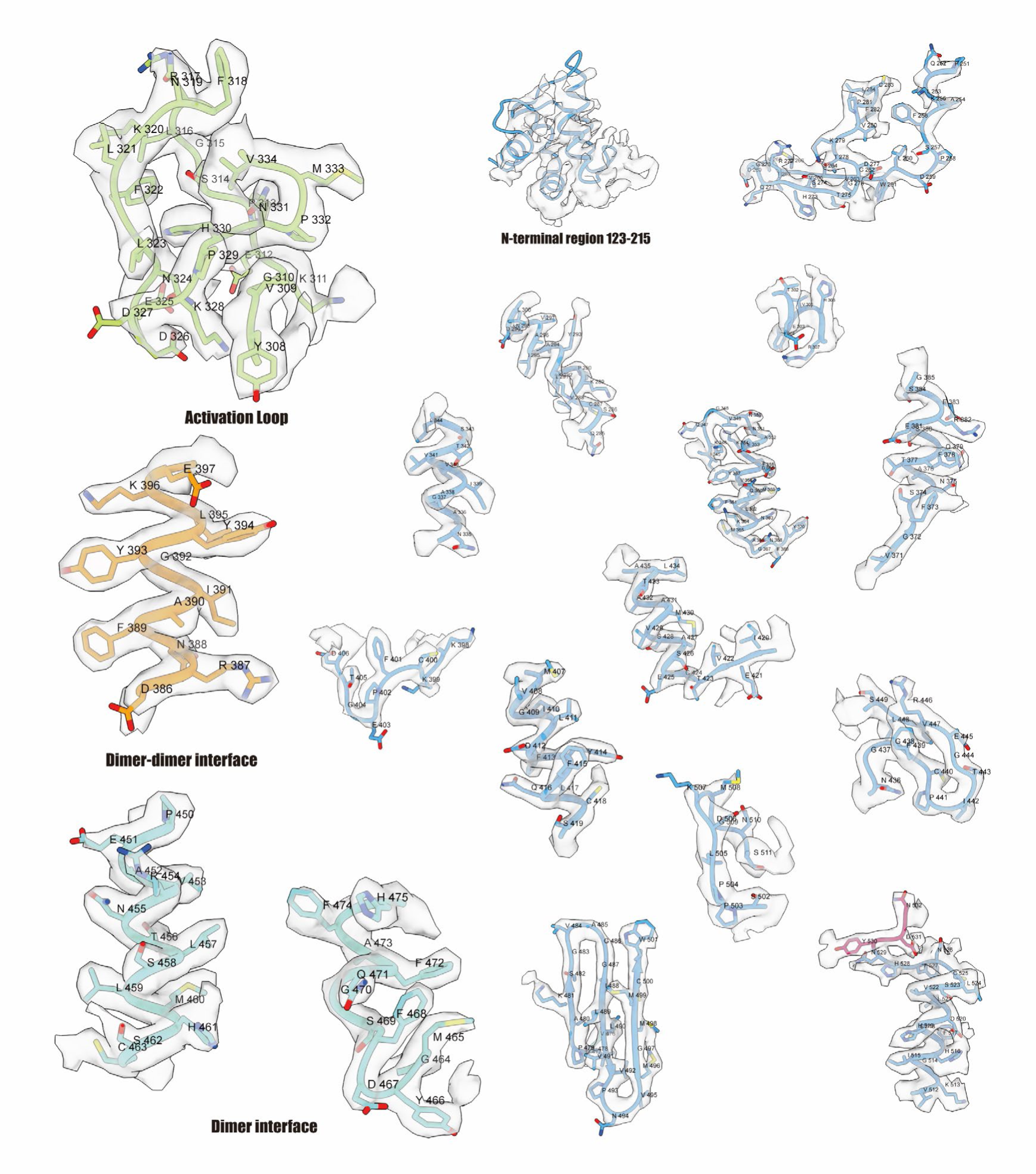
Representative EM detail. The transparent surface represents the density map, and the amino acid residues are indicated with single-letter abbreviations.

**Fig. S5.**
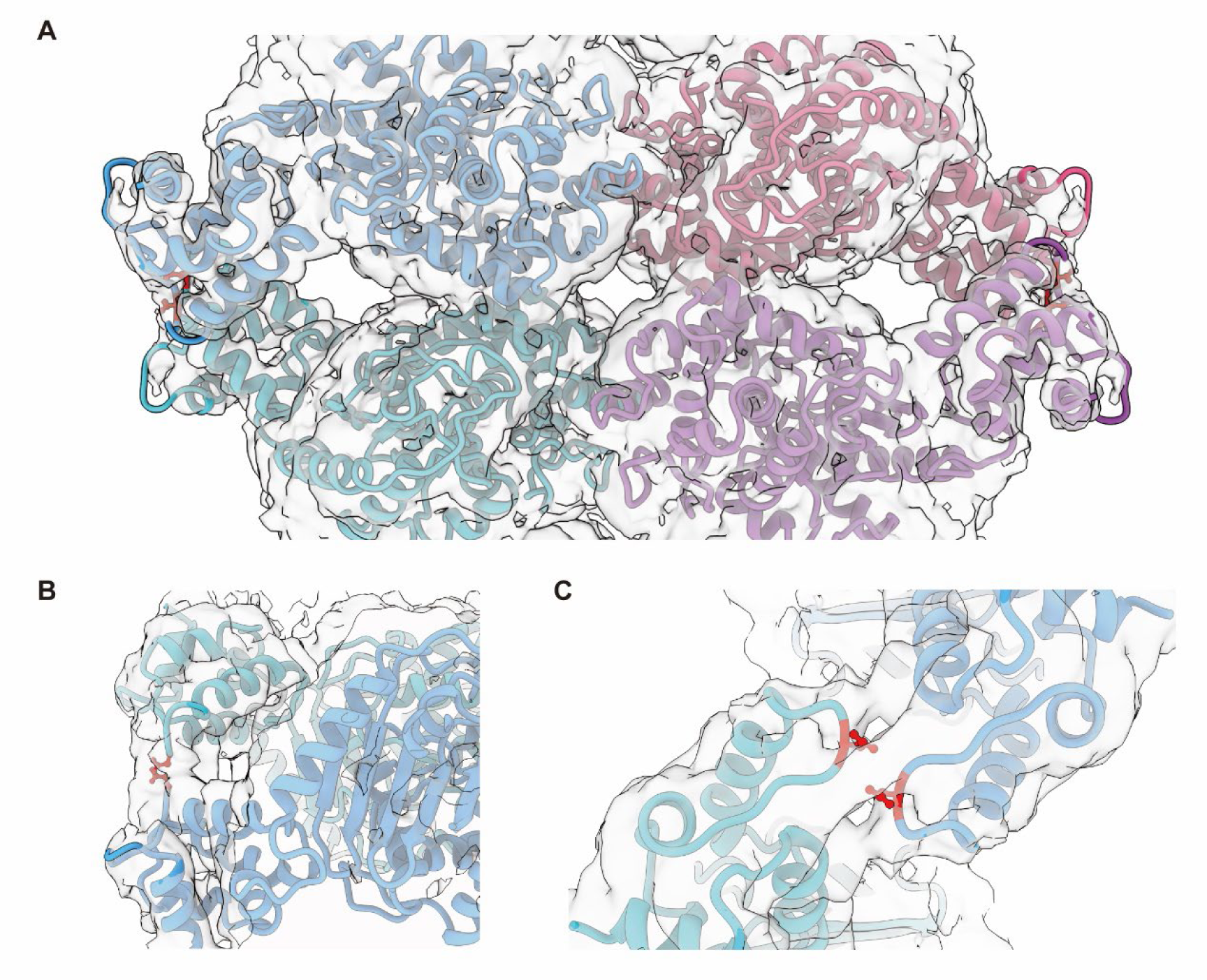
Filamentation interaction of GAC-Pi in the N-terminal region. A) Front view of the helical interface of GAC-Pi with density map and the possible residues involved in the interaction are colored in red. B) Top view of the helical interface. C) Side view of the helical interface.

**Fig. S6.**
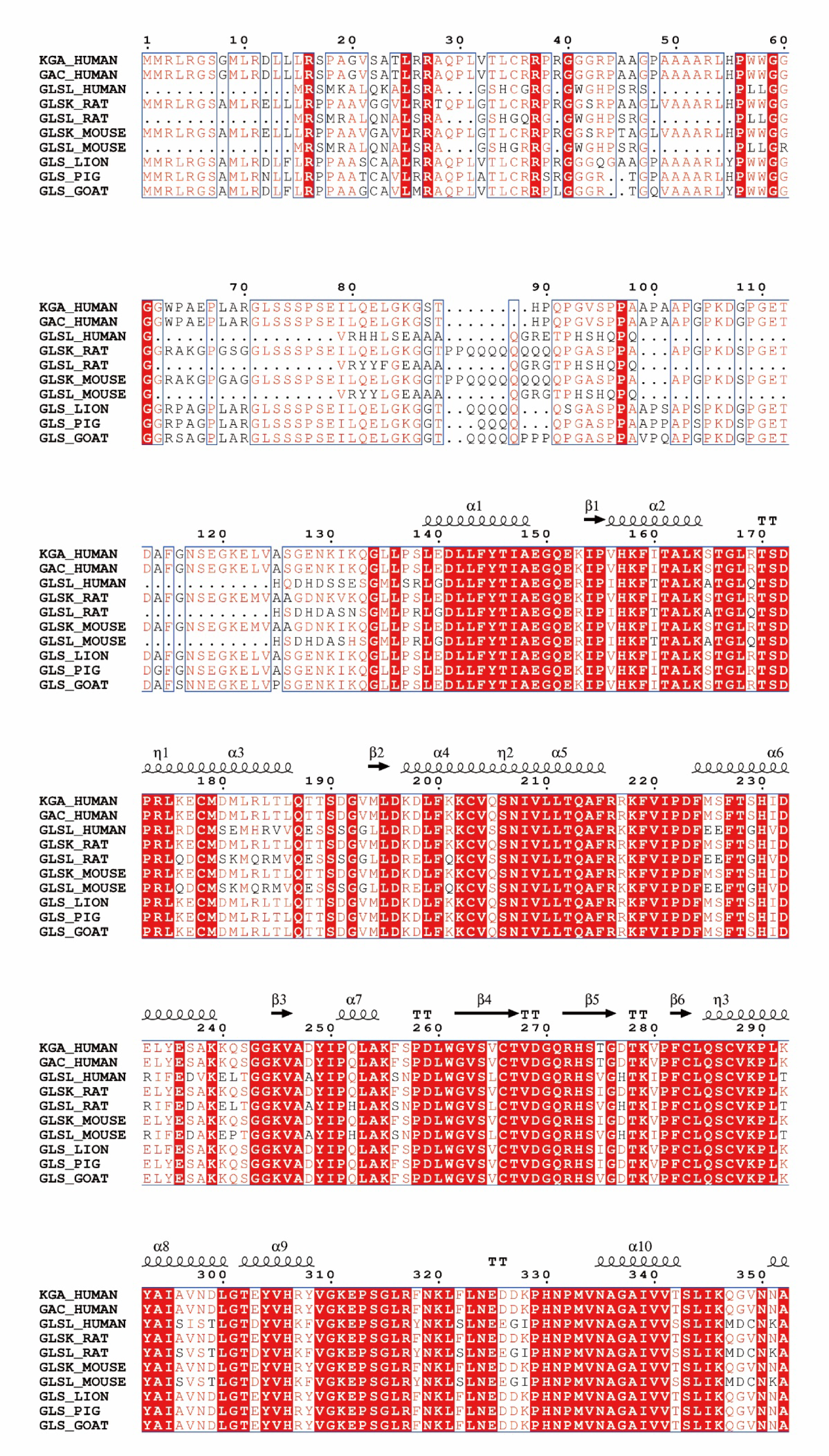
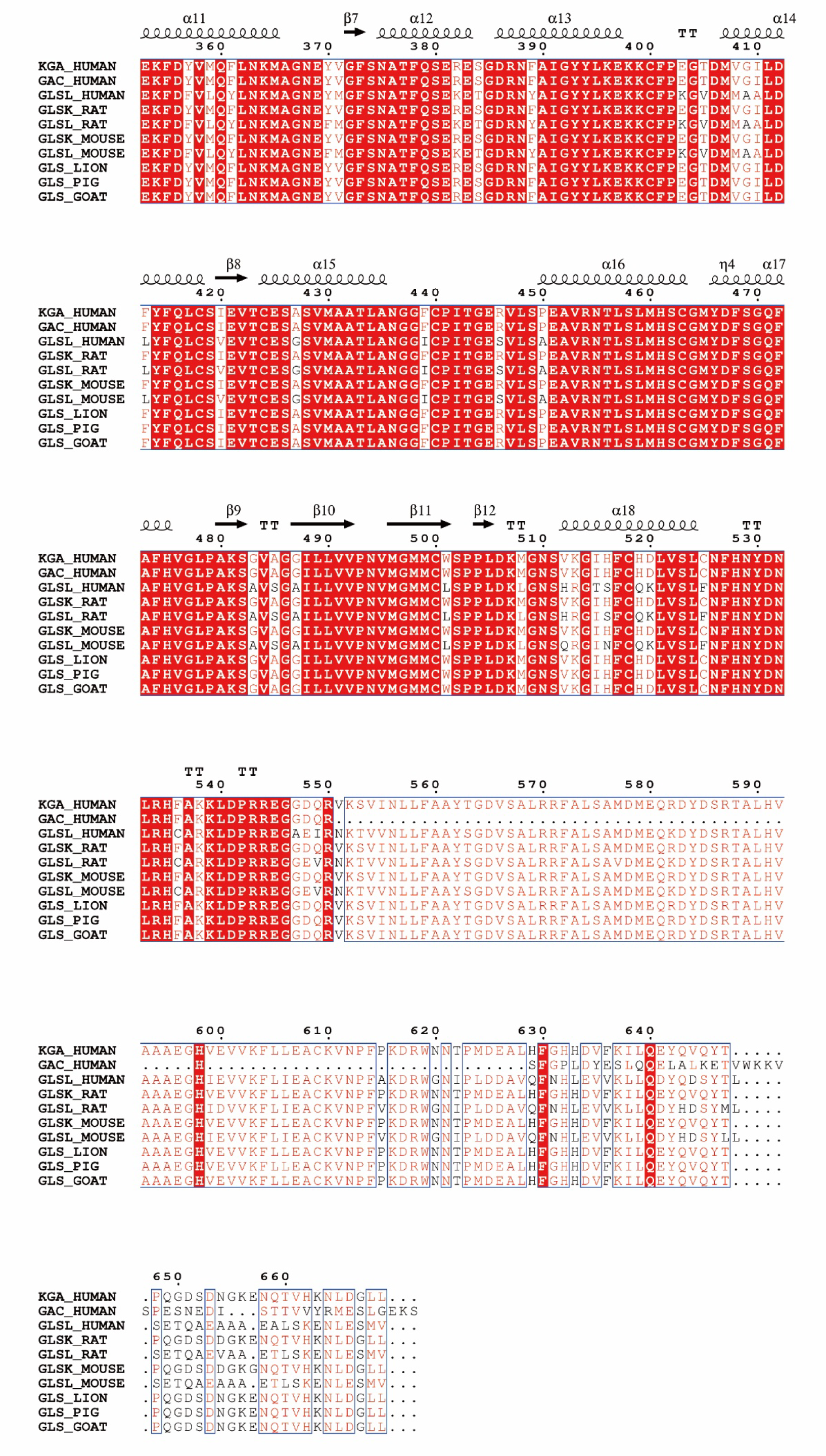
Seqence alignment of GLS from different mammalian species. The sequence alignment of the amino acid sequence of human KGA and GAC, human GLSL, rat GLSK, rat GLSL, mouse GLSK, mouse GLSL, lion GLS, pig GLS, and goat GLS is shown. The conserved residues are shaded in red, and secondary structure information is indicated above the alignment.

**Fig. S7.**
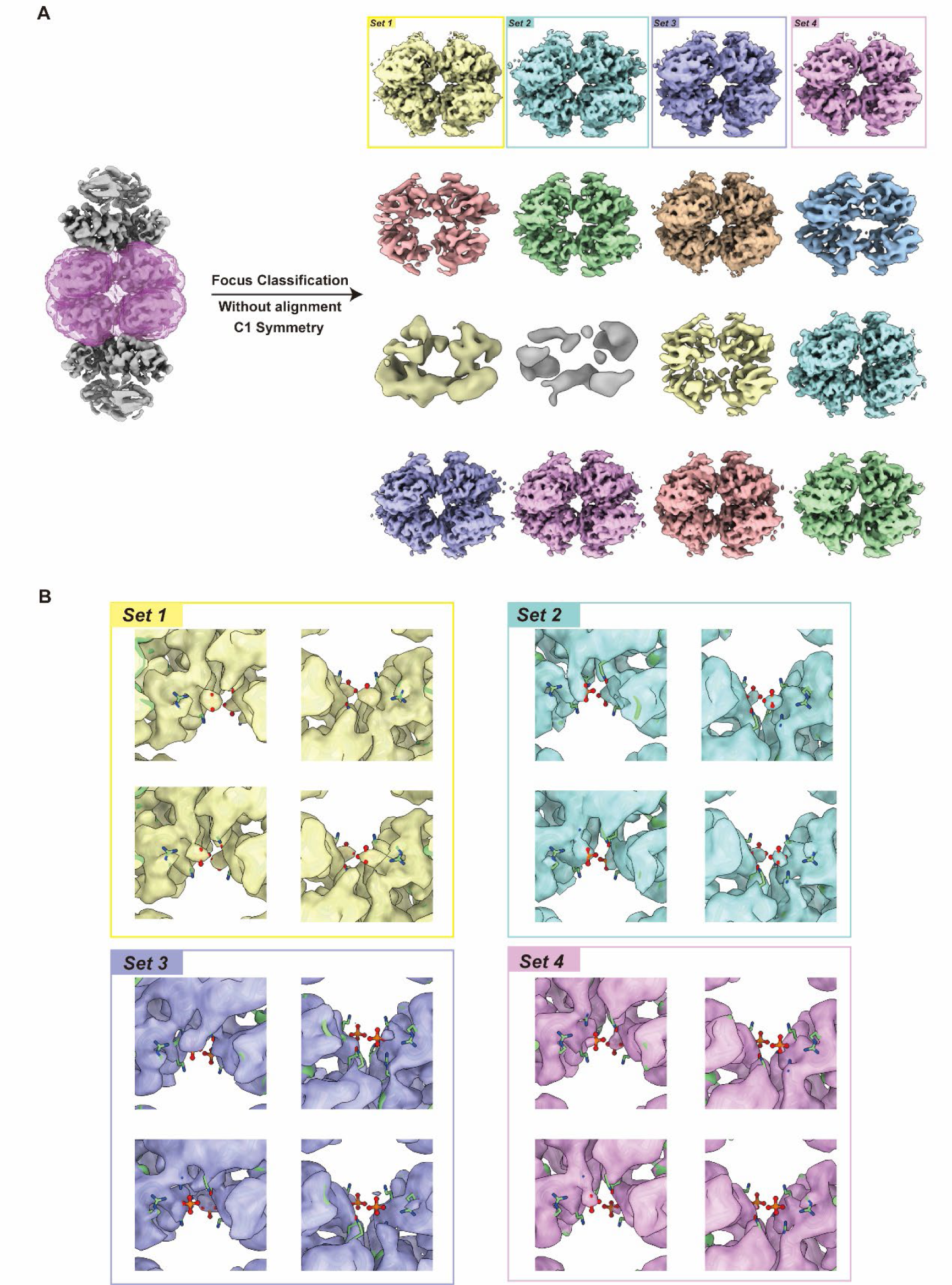
Flexibility of the Pi binding pocket. A) 16 classes generated by 3D classification without alignment using C1 symmetry with a mask focusing on the catalytic core. B) Zoom-in of the Pi binding pocket for 4 selected classes above. And the model of GAC-PF is shown as a reference.

**Fig. S8.**
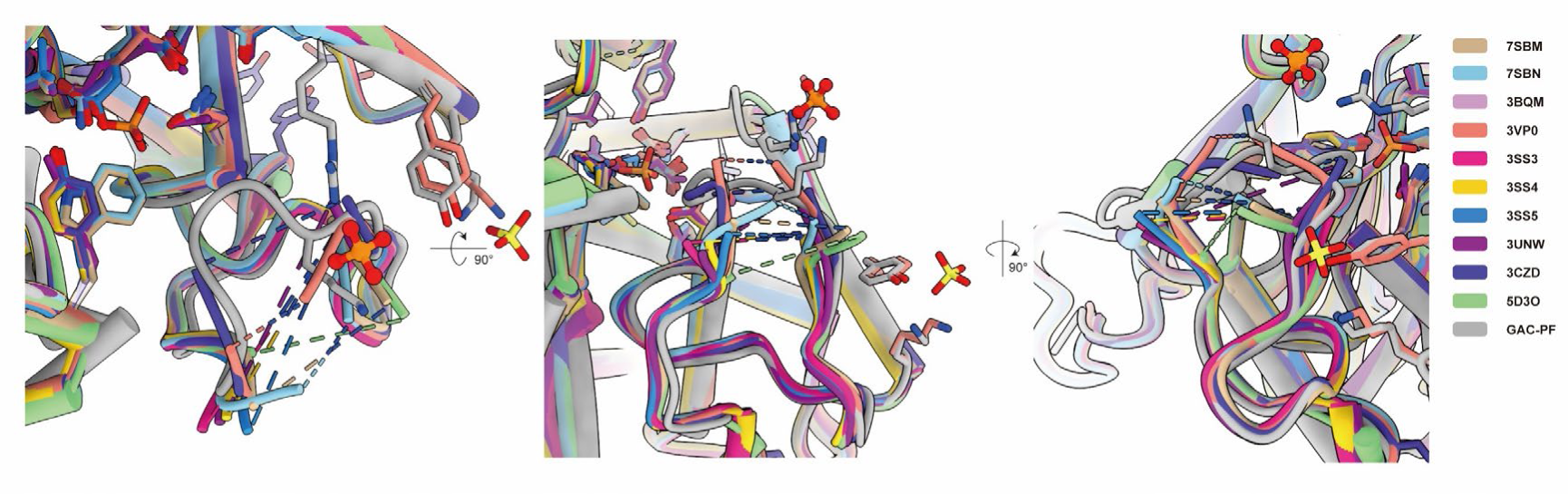
High flexibility of AL in mammalian glutaminase model without inhibitor binding. Comparison of the AL in GAC-PF with different models. The disordered loop is indicated by dashes. The AL in other models is shown to be highly flexible, suggesting its potential role in regulating the enzyme activity.

**Fig. S9.**
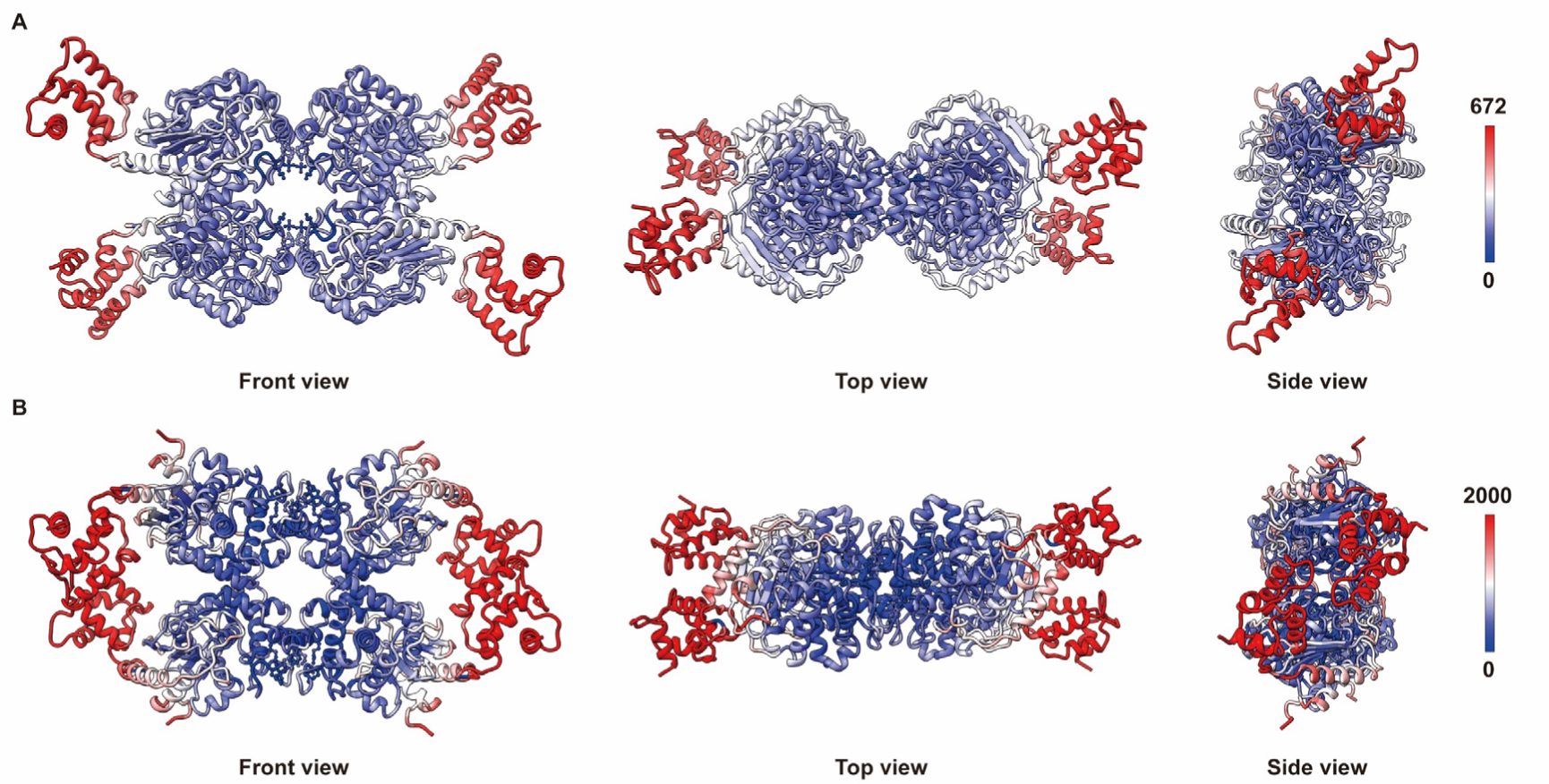
B-factor analysis of GAC-PF model. **A)** B-factor analysis of the helical unit model, with the color gradient representing the B-factor values ranging from low (blue) to high (red). **B)** B-factor analysis of the helical interface, with the color gradient representing the B-factor values ranging from low (blue) to high (red).

**Fig. S10.**
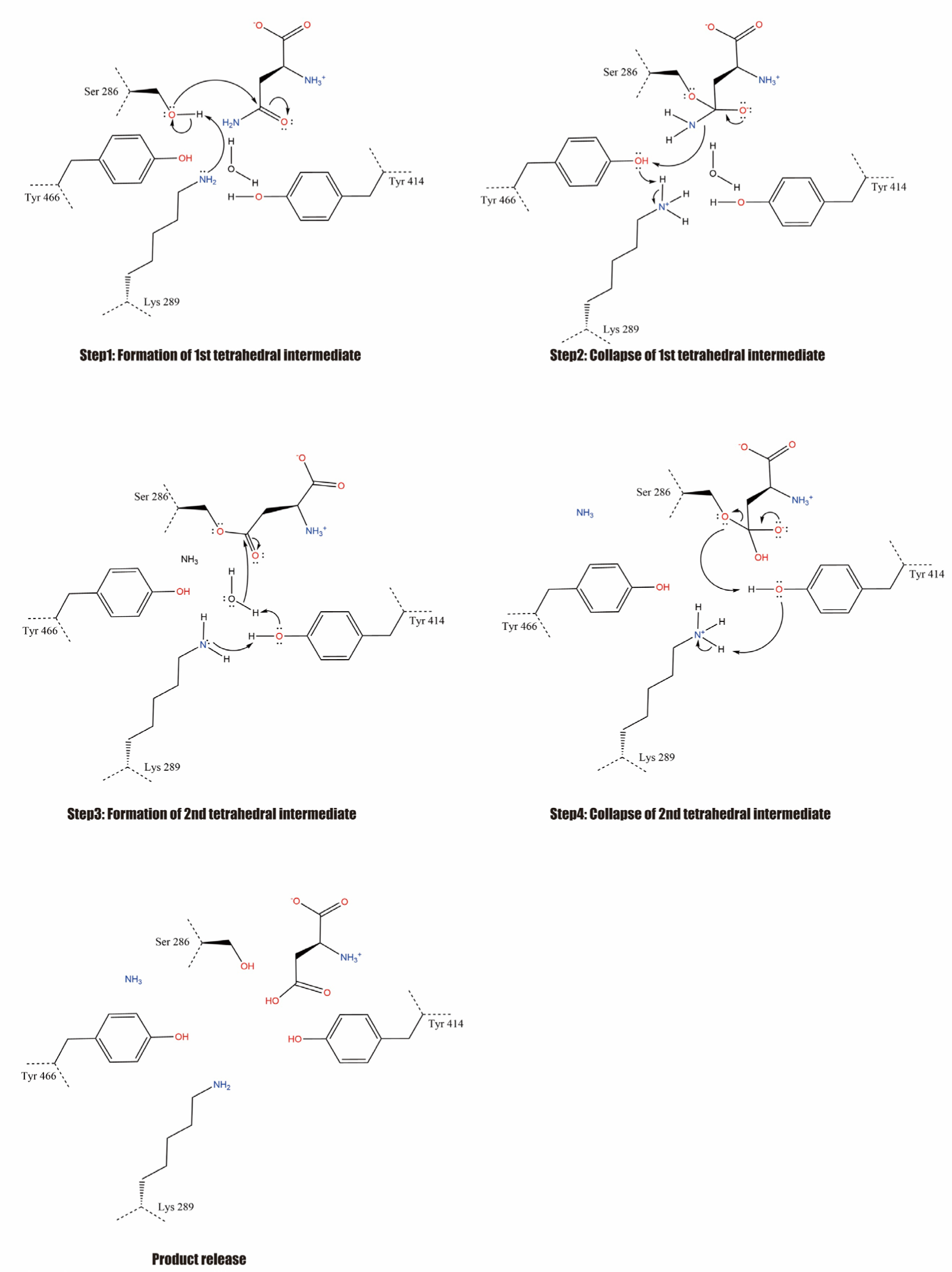
Proposed catalytic diagram for glutaminase. The catalytic reaction of GAC can be roughly divided into four steps: 1) K289 deprotonates S286, initiating a nucleophilic addition onto the carbonyl carbon of the substrate amide group, forming the first tetrahedral intermediate; 2) The first tetrahedral intermediate collapses, cleaving the C-N bond and releasing the ammonia product, the nitrogen of which deprotonates K289 via a proton relay through Y466; 3) K289 deprotonates water via a proton relay with Y414, initiating a nucleophilic addition at the carbonyl carbon, forming a new tetrahedral intermediate; 4) The new tetrahedral intermediate collapses, cleaving the acyl-enzyme bond and liberating S286, which in turn deprotonates K289 via a proton relay with Y414.

**Fig. S11.**
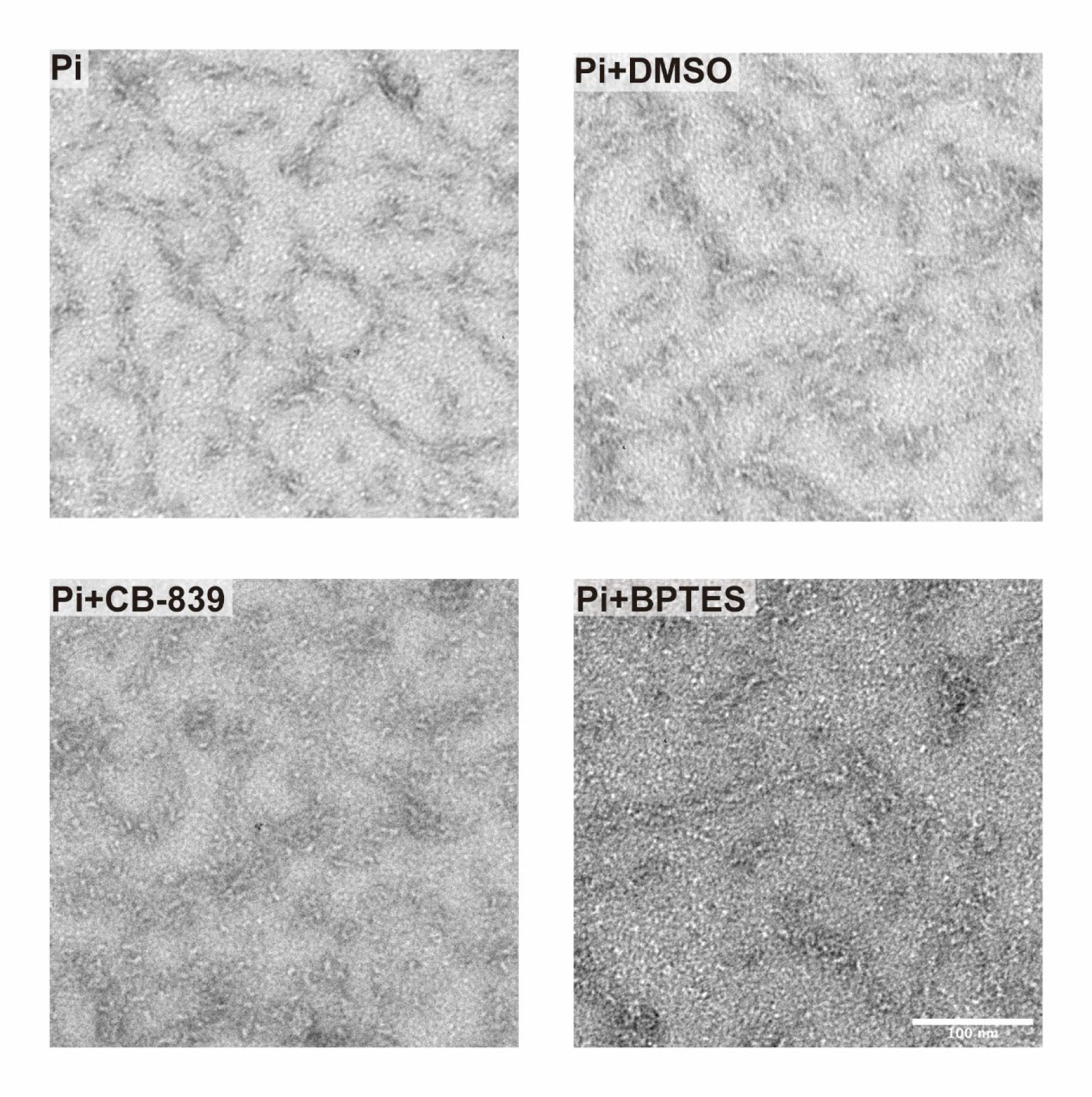
Negative staining images for inhibitor joined sample. Negative stain electron microscopy micrographs of GAC^Pi^ with or without inhibitors. Fewer filaments were observed after incubating with inhibitors BPTES or CB-839, with DMSO as a negative control. Scale bar represents 100 nm.

**Fig. S12.**
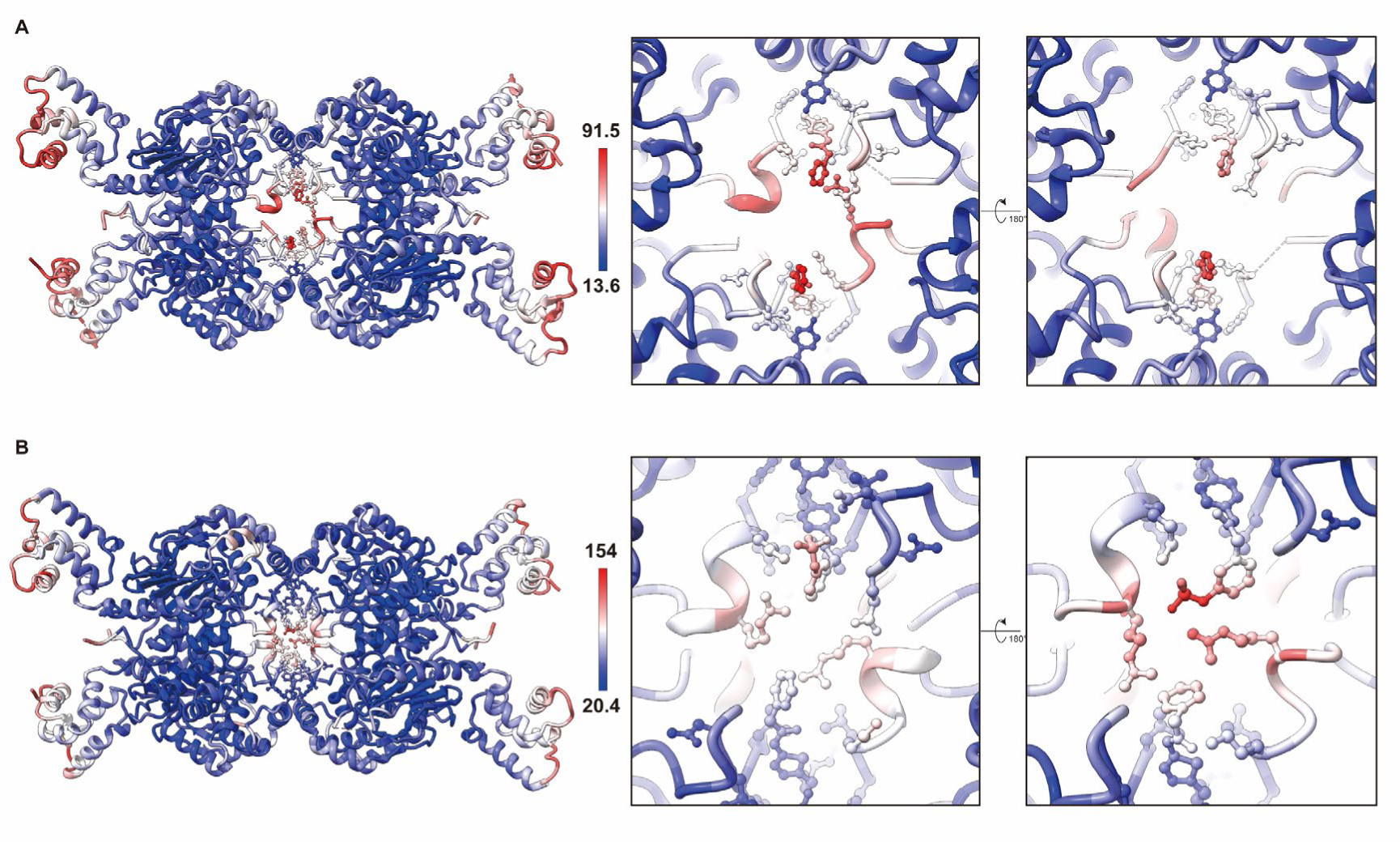
B-factor analysis of BPTES and CB-839 binding to the allosteric site in the GAC crystal structures. A) The model of GAC in complex with BPTES (PDB ID:3UO9). The terminal of BPTES shows relative flexibility, and AL loop is not fully fixed. B) The model of GAC in complex with CB-839 (PDB ID:5HL1).

**Supplementary Table S1.** Cryo-EM data collection and model refinement.

|  | helical interface<br>(EMD-35574, PDB 8IMB) | helical unit<br>(EMD-35573, PDB 8IMA) |
| --- | --- | --- |
| **Data collection** |  |  |
| EM equipment | Titan Krios | Titan Krios |
| Detector | K3 camera | K3 camera |
| Magnification | 22,500x | 22,500x |
| Voltage (kV) | 300 | 300 |
| Electron exposure ((e-/Å<sup>2</sup>)) | 50 | 50 |
| Defocus range(μm) | -1.2 to -1.8 | -1.2 to -1.8 |
| Pixel size(Å) | 1.06 | 1.06 |
| Symmetry imposed | D2 | D2 |
| Number of collected movies | 9265 | 9265 |
| Initial particle images (no.) | 3271231 | 3271231 |
| Final particle images (no.) | 321414 | 203004 |
| Map resolution (Å) | 2.9 | 2.9 |
| FSC threshold | 0.143 | 0.143 |
| Map resolution range (Å) | 2.8-3.8 | 2.8-3.7 |
| **Refinement** |  |  |
| Initial model used (PDB code) | 3SS4 | 3SS4 |
| Map sharpening B-factor(Å<sup>2</sup>) | 80 | 80 |
| Model composition |  |  |
| Non-hydrogen atoms | 12304 | 12304 |
| Protein residues | 1580 | 1580 |
| Ligands | PO4: 4 | PO4: 4 |
| Waters | 0 | 0 |
| B factors(Å<sup>2</sup>) | 80 | 80 |
| R.m.s. deviations |  |  |
| Bond lengths (Å) | 0.004 | 0.004 |
| Bond angles (°) | 0.564 | 0.54 |
| Validation |  |  |
| MolProbity score | 1.83 | 2.37 |
| Clashscore | 4.25 | 9.97 |
| Poor rotamers (%) | 1.75 | 2.92 |
| Ramachandran plot |  |  |
| Favored (%) | 93.19 | 91.79 |
| Allowed (%) | 6.81 | 8.21 |
| Disallowed (%) | 0 | 0 |

